# Using Packing Defects in Heterogeneous Biological Membrane as a Lens to Explore Protein Localization Propensity and Small Molecule Permeability

**DOI:** 10.1101/2022.09.20.508633

**Authors:** Madhusmita Tripathy, Anand Srivastava

## Abstract

Plasma membrane (PM) heterogeneity has long been implicated in various cellular functions. However, mechanistic principles governing functional regulations of lipid environment is not well understood due to the inherent complexities associated with the relevant length and time scales that limit both direct experimental measurements and their interpretation. In this context, computer simulation holds immense potential to investigate molecular-level interactions that lead to PM heterogeneity and the related functions. Herein, we investigate spatial and dynamic heterogeneity in model membranes with coexisting liquid ordered and liquid disordered phases and characterize the membrane order in terms of the topological changes in lipid local environment using the non-affine parameter (NAP) framework. Furthermore, we probe the packing defects in membrane with coexisting fluid phases, which can be considered as the conjugate of membrane order assessed in terms of the NAP. In doing so, we formalize the connection between membrane packing and local membrane order and use that to explore the mechanistic principles behind preferential localization of proteins in mixed phase membranes and membrane permeability of small molecules. Our observations suggest that heterogeneity in mixed phase membranes follow some generic features, where functions may arise based on packing-related basic design principles.

**Significance:** Functionally important complex lateral and transverse structures in biological membrane result from the differential molecular interactions among a rich variety of lipids and other building blocks. The nature of molecular packing in membrane is a manifestation of these interactions. In this work, using some of the ideas from the Physics of amorphous materials and glasses, we quantify the correlation between heterogeneous membrane organization and the three dimensional packing defects. Subsequently, we investigate the packing-based molecular design-level features that drive preferential localization of peptides in heterogeneous membrane and membrane permeation of small molecules.

## INTRODUCTION

About 150 years ago, from their simple but ingenious experiments on movement of anesthetic and other small solutes between the surrounding and cell interior, Charles Ernest Overton and Hans Horst Meyer made the astute observation that the cell boundaries could be composed of “aliphatic” lipid molecules^1,2^. The seminal work of Overton and Meyer established the idea of biological membrane as a semi-permeable boundary and presumably laid the foundation for membrane biophysics. The ensuing early activities towards understanding the structure and organization of the biological membrane from the likes of Pockels and Langmuir^3^, Gortner and Grendel^4^ and Davison and Danielli^5^ led to the now well-known information that biological membrane are made up of two layers of lipid leaflets that are stacked such that the hydrophobic fatty-acid tails make the core of the bilayer and the polar head groups face the aqueous media on both sides. The paradigms of membrane organization has undergone a cascade of changes since then^6,7^ with a large body of evidence pointing towards a complex lateral organization with existence of non-random localization of lipids and proteins on membrane surface resulting in existence of physiologically functional heterogeneous sub-100 nm patterns that we now call “rafts” in living membranes^8–14^. With the advent of single molecule tracking techniques and super resolution microscopy and tomography^15–18^ as well as lipidomics data^19–27^, we now have access to unprecedented details on membrane composition, structure and dynamics.

Over the last few decades, spatial and dynamic heterogeneity in membrane has been systematically and extensively investigated, in both reconstituted model membranes at carefully chosen composition and cell derived/living cell membranes ^8,10,28–39^. However, our understanding of their functional implications still remains far from complete, mostly due to the lack of a comprehensive molecular level picture. We point to a few reviews that have succinctly laid down the advances made in the field and also highlighted the path forward towards more clearly elucidating the structure, dynamics and functional role of the membrane “rafts”^40–46^. Traditionally, lipid rafts have been identified as the heterogeneous, highly dynamic, cholesterol and sphingomyelin rich tightly packed domains that are implicated in specific protein recruitment. In model membranes, the liquid ordered (*L*_*o*_) domains in a phase separated membrane are considered to be the prototype for rafts ^39,47,48^ with tighter packing and slower diffusion as compared to a more fluid liquid disordered (*L*_*d*_) regions. In general, with diffusion and packing behavior as distinguishing features of the two co-existing fluid phases in membrane systems, the relationship between the molecular packing of lipids, mobility and membrane order is seemingly obvious from the classical free-volume based theories of diffusion^49,50^. However, despite the tremendous advances made in experimental resolutions in both time and space, accurate quantification and formulation of this correlation is still elusive^51–55^. In this regard, computer simulation has proven to be a very useful tool and the success of ‘computational microscopy’^56,57^ spans from the ability to observe phase separated nano-structures, reminiscent of the ordered raft domains, in model binary/ternary lipid systems ^47,58,59^ to investigating their plausible functional implications^60,61^. Although our understanding of these rafts and their functions is limited to such simplified model membranes, computer simulations have been able to provide further insights into their microscopic structure and function. Only recently, relatively simple model membranes exhibiting liquid ordered (*L*_*o*_) and disordered (*L*_*d*_) phases have been able to elucidate the distinct nature of lipid packing within the *L*_*o*_ domains^59,62,63^, which are observed to play a crucial role in membrane permeation of small molecules^64–66^.

Lipid packing defects^67^, the transiently exposed hydrophobic tails of the lipids, are the cooperative manifestation of both the membrane order and packing (which results from differential lipid interactions), together with the collective dynamics of the lipids in the ordered/disordered domains. It can, therefore, be loosely considered as the conjugate of local membrane orderliness. While non-trivial to identify in experiments, packing defects has been extensively studied in computer simulation^60,68–71^. Over the last decade, the nature of packing defects in curved membranes have been well characterized in various simulation studies, unraveling their role in the adsorption of various proteins that contain specific amphipathic helices^60,69,70,72^. Amphipathic lipid packing sensor (ALPS) motifs in these proteins, can sense large pre-existing packing defects and initiate binding by anchoring some of their bulky amino acid residues into such defects. Such studies on the curvature dependent binding of ALPS motifs have been able to elucidate the binding mechanism of various peripheral and cytosolic proteins and their physiological implications in cells^73–77^. However, a similar understanding of the nature of lipid packing defects in coexisting fluid phase membranes is still missing, even though the lateral organization in a realistic cell membranes has highly heterogeneous characteristics. One important physiological function of the raft-like domains therein is the partitioning of peripheral proteins^78–80^, where large packing defects are known to provide suitable platforms for membrane adsorption^70,81^. Various experiments have shown preferential domain affinities for distinct membrane binding motifs to either of the *L*_o_ or *L*_d_ domains or their interface. For example, while most of the peripheral proteins such as RAB proteins RAB1, RAB5, and RAB6 preferentially bind to *L*_d_ domains^82^, HIV gp41 interacts predominantly at the *L*_o_*/L*_d_ domain boundary^83^ and aspirin (acetylsalicylic acid) binds to *L*_o_ domains^84^. Intriguingly, the ordered domain affinity has been found to be further governed by the relative contrast between the ordered and disordered domains, which is known to be rather subtle in cell derived giant plasma membrane vesicles (GPMVs) and quite distinct in giant unilamellar vesicles (GUVs)^37^. Accordingly, the palmitoylated trans-membrane domain of Linker for Activation of T Cells (tLAT) is shown to bind to the *L*_o_ domains in GPMV^85,86^, while they are specifically excluded from *L*_o_ phase in GUVs^87^. While several other proteins exhibit such selective domain preference^80^, the basic design principle behind this specificity of membrane-protein interactions in mixed phase membrane is so far unexplored.

Interestingly, the nature of packing defects in pure phase (homogeneous) membranes are found to be rather generic, as they exhibit a few characteristic features that depend only on the overall order (phase) of the membrane^71^. In general, pure *L*_o_ systems are relatively scarce in packing defects, while pure *L*_d_ systems exhibit a denser distribution of defects. Furthermore, the defects in the *L*_o_ systems are observed to be significantly smaller (and shallower) as compared to those in the *L*_d_ systems. While these observations can be trivially extrapolated to coexisting (mixed phase) *L*_o_*/L*_d_ membranes, wherein the *L*_*d*_ regions are dominated in terms of packing defects^71,88^, rationalizing their role in peripheral protein partitioning is nontrivial, as proteins *do not* exclusively partition with the *L*_d_ domains that are rich in large and deep defects. So, what factors decide the domain affinity in such membranes?

Beyond protein partitioning, lipid ordering can also play a crucial role in the membrane permeability of small molecules. A recent study by Richard Pastor and co-workers ^89^ investigated the transport of oxygen and water molecules through ternary membrane systems exhibiting homogeneous *L*_*o*_ and *L*_*d*_ phases. Their observations indicate the existence of separate permeation pathways in *L*_*o*_ and *L*_*d*_ phase membranes, which have their origins in specific packing of of lipids. They showed a nearly free diffusion in bilayer mid-plane in *L*_*o*_ membranes while the *L*_*d*_ membranes were statistically more permeable. Their observations on relative concentration of oxygen in the bilayer mid-plane of both membranes were further supported by electron paramagnetic resonance spectroscopy. It is therefore evident that membrane packing and order not only dictate the peripheral membrane protein association and localization, but also the trans-bilayer permeation pathways. Traditionally, lateral lipid diffusion in experiments has been modeled in terms of free volume theories^90^, which in its simplest form assumes that the dynamics of molecules in liquids and glasses proceed in discrete diffusive jumps and critically depend on the available free volume^91^. In the case of lipid membrane, this free volume can be related to the fluctuation in area per lipid. However, unlike the case of molecular liquids, this free volume with complex lipid geometry can not be defined unambiguously^92^. In spite of several rigorous attempts in the past, a complete understanding on the role of free volume in lipid diffusion still remains missing^68,92–95^. The presence of multiple lipid species and the intrinsic heterogeneity of the membrane further complicates the problem. Moreover, processes such as membrane permeability, protein-membrane interaction, signalling across the membrane and other functions, which closely depend on inter-leaflet coupling and involve both lateral and transverse diffusion, demands explanations that go beyond the simple free volume theories.

In this study, we investigate the functional significance of membrane orderliness in the light of packing defects. Our goal is to understand and formalize the correlation between the membrane order in coexisting (mixed phase) *L*_o_*/L*_d_ membranes and the packing defects therein, where pure phase membranes serve as control systems for such investigations. Towards this, we characterize the local membrane orderliness in terms of the non-affine parameter (NAP)^63,88^, which captures the distinct nature of spatio-temporal evolution of lipids in their local neighbourhoods without any knowledge on lipid chemistry. We identify the threedimensional packing defects in these membranes using our recently developed algorithm^71^, which efficiently circumvents the computational bottleneck of grid based calculations. Our results indicate a direct correlation between orderliness and defects for three different mixed phase membrane systems with distinct lipid constituents, suggesting the correlation to be rather generic. Moreover, a stronger correlation indicates the nature of mixed phase membrane to be more ordered-like, in which case, protein can also partition onto the ordered domains. Furthermore, the dynamics of these defects in the *L*_o_ and *L*_d_ domains are observed to be distinctly different, indicating two possible scenarios for protein partitioning onto any mixed phase membranes. Using the framework of NAP, we further investigate the membrane partitioning of tLAT and find that palmitoylation indeed increases its ordered phase affinity. By analyzing the surface defects (hydrophobic defect pockets) and core defects (free volume in the membrane mid-plane) in the membrane systems studied by Ghysels et al.^89^, we find that while the *L*_*d*_ membrane clearly dominates in terms of size of surface defects, the trend is reversed for core defects, i.e., *L*_*o*_ systems exhibit larger free volume in the membrane core. This can provide a valid explanation on the free transverse diffusion in *L*_*o*_ domains, while *L*_*d*_ domains remain statistically more permeable. The membrane packing and order can, thus, influence both its peripheral and trans-bilayer functions. Our inferences are rather general and can be extended to any mixed phase membrane irrespective of its chemical nature.

## RESULTS

### Local orderliness dictates the defect profiles in mixed phase membranes

The nature of packing defects in mixed phase membranes is expected to be intrinsically coupled to the membrane order. The membrane order is traditionally characterized in terms of the local lipid arrangements, membrane packing (area per lipid), thickness, and lipid tail-order parameter(S_CD_)^59,62,63^. However, the dynamic nature of these regions, wherein lipids constantly enter and exit them, necessitate the inclusion of temporal information in the analysis. Herein, we quantify the orderliness of three mixed phase ternary lipid systems, DPPC/DOPC/CHOL, PSM/DOPC/CHOL, and PSM/POPC/CHOL membranes, in terms of the non-affine parameter (NAP), i.e., the residual non-affine content of the deformation (χ^2^)^96,97^, which can capture the distinct nature of the spatio-temporal evolution of lipids in their local neighborhood^63^ as shown in Fig. 1a (Please see Methods sections for details on NAP calculations). Furthermore, this analysis can also identify local molecular scale heterogeneity in pure phase systems indicating regions that undergo more heterogeneous topological rearrangements than their neighborhoods^63,88^, and thus, is a sensitive marker of membrane order.

**FIG. 1:**
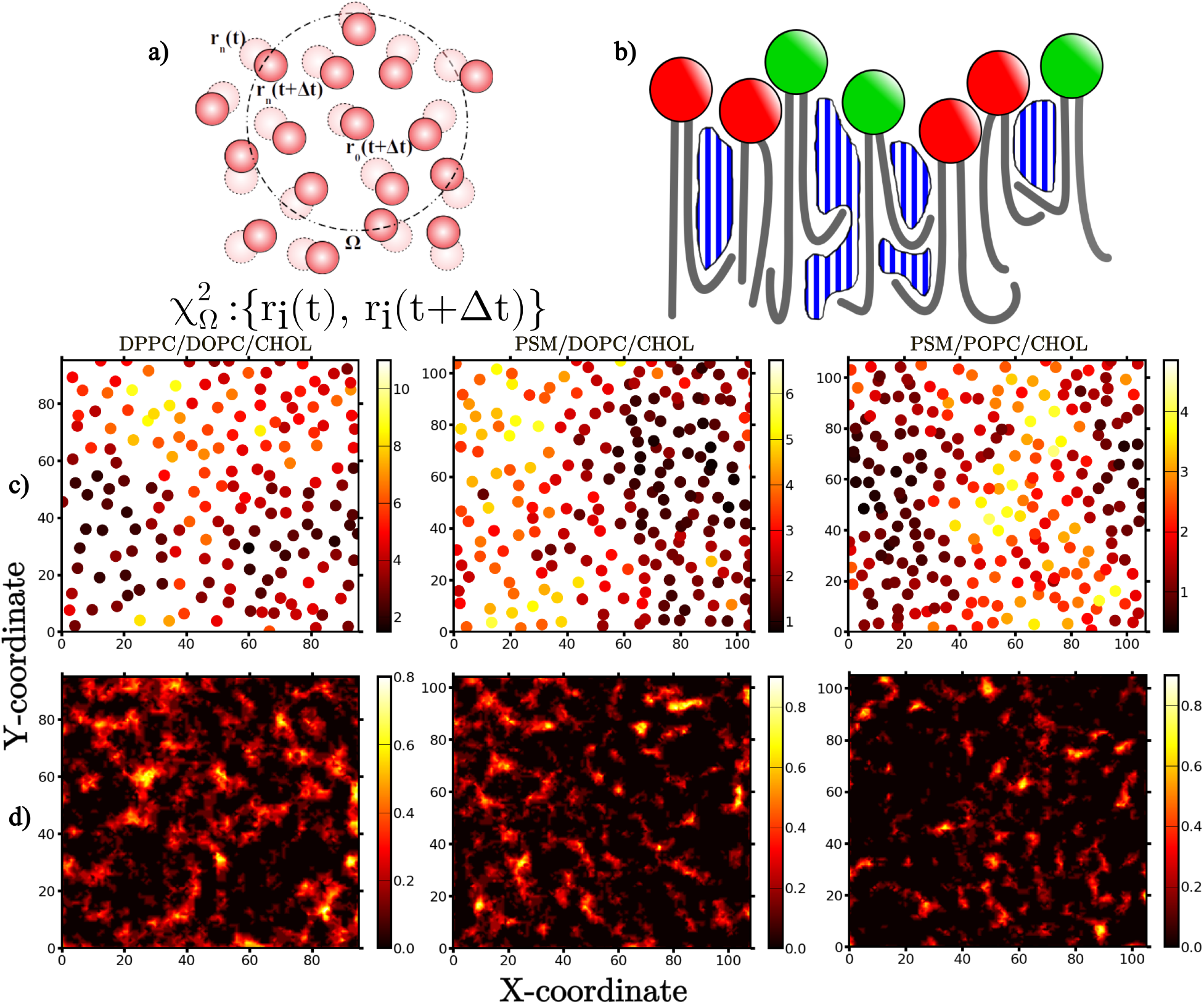
Spatial correlation of χ^2^ values and packing defects. a) The definition of neighborhood Ω in the χ^2^ analysis. *r*_*i*_ represents the coordinates of reference lipid sites, whose spatial and temporal evolution is used to calculate χ^2^. b) Schematic indicating the various defects that can be identified using the 3-dimensional defect analysis, shown as blue/while shaded regions. Note that the lipid reference sites are specifically excluded while identifying the defects c) χ^2^ spatial map and d) defect spatial map for DPPC/DOPC/CHOL (left), PSM/DOPC/CHOL (middle), and PSM/POPC/CHOL (right) systems, each exhibiting mixed *L*_*o*_*/L*_*d*_ phases.

Our earlier works on pure phase systems have indicated that irrespective of lipid chemistry, lipids in pure *L*_o_ systems exhibit consistently low χ^2^ values as compared to those in pure *L*_d_ systems^63,88^. In mixed phase membranes, the local ordering of lipids can be distinctly different, which can subsequently result in specific packing defects profiles. To understand this, we compare the χ^2^ maps, against the spatial distribution maps of the defects (see Methods) in the three membranes as shown in Fig. 1. We observe that regions in these membranes that possess low χ^2^ value, i.e., the ordered domains, are mostly defect-free. Similarly, a large defect can almost always be mapped to reference sites with high χ^2^ values, i,e., disordered domains. These observations are valid for all the three systems and thus, can be a general feature of all mixed phase systems. It, therefore, appears that the observations on the nature of defects in pure phase membranes can be extended to mixed phase membranes: ordered domains in mixed phase membrane remain relatively defect-free as compared to disordered domains.

However, as shown in Figure 2, the distributions of defect size for the three mixed phase systems are rather distinctive when compared to their pure phase membrane counterparts: in the case of DPPC/DOPC/CHOL, the distribution is very similar to its *L*_d_ counterpart, for PSM/POPC/CHOL system it is comparable to the corresponding *L*_o_ one, and for PSM/DOPC/CHOL system, it is found to be intermediate to the corresponding two pure systems. Thus, the mixed phase membranes lack a generic trend in terms of the defect size, as compared to the pure phase membranes, wherein the *L*_o_ membranes always exhibit smaller defects as compared to their *L*_d_ counterparts. The origin of this observation can be again related to the orderliness of the membrane, characterized in terms of the χ^2^ distribution. Pure phase *L*_o_ systems always exhibit a sharper and narrower distributions of χ^2^ values as compared to the corresponding *L*_d_ ones, which is a generic feature of pure phase (homogeneous) membranes^63,71,88^. Interestingly, χ^2^ distribution for the three mixed phase systems follow the exact same trend as the defect size distributions, indicating the mixed phase DPPC/DOPC/CHOL membrane to be more disordered-like and mixed phase PSM/POPC/CHOL membrane to be more ordered-like. The mixed phase PSM/DOPC/CHOL membrane is found to be intermediate to the two pure systems.

**FIG. 2:**
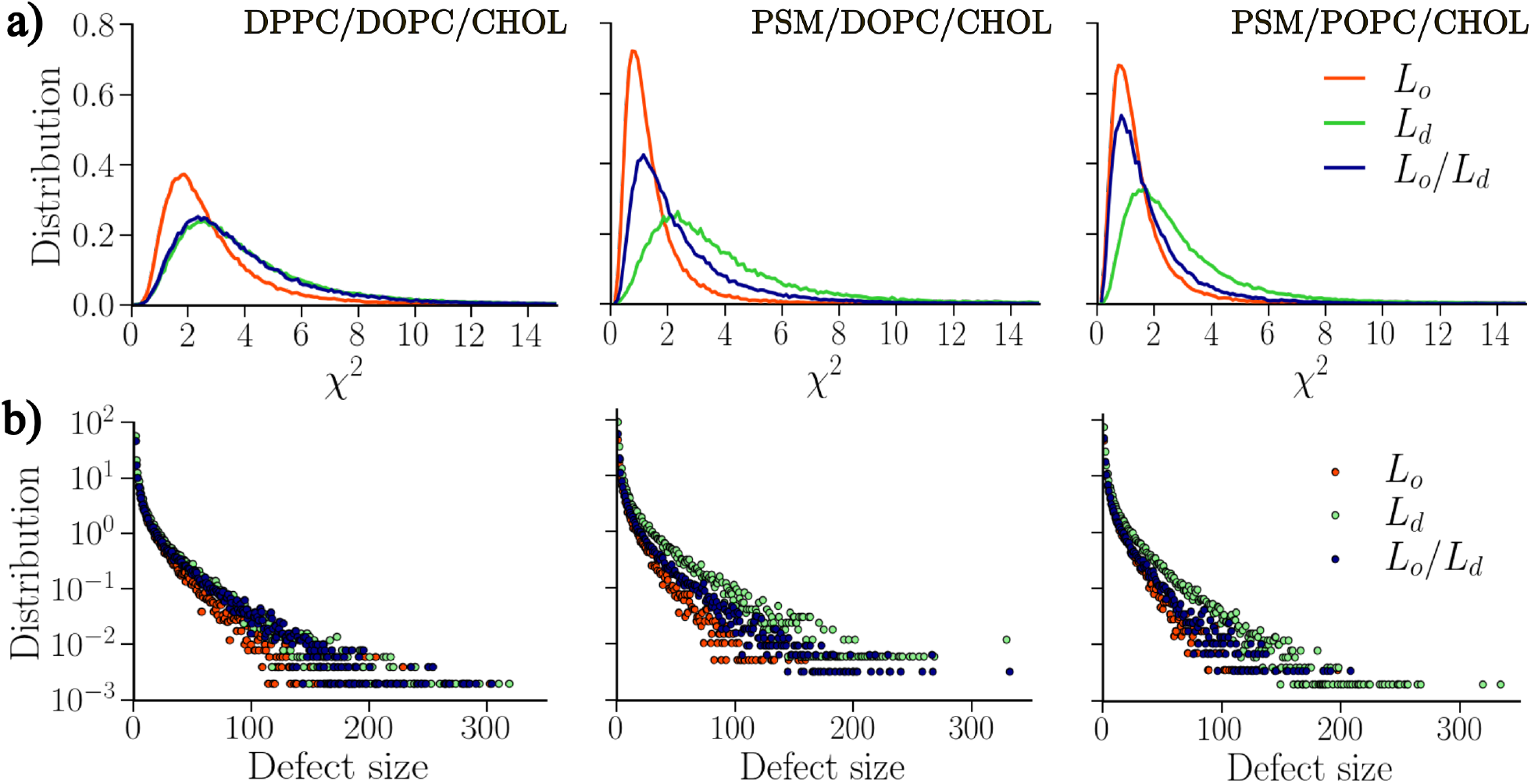
Overall correlation between the χ^2^ values and defects. Distributions of (a) χ^2^ values and (b) defect size for DPPC/DOPC/CHOL (left), PSM/DOPC/CHOL (middle), and PSM/POPC/CHOL (right) systems exhibiting pure and mixed liquid phases. The range of axes are kept the same for better comparison.

This apparent correlation between χ^2^ values and the packing defects can be formalized in terms of a two dimensional probability distribution P(*n*_*d*_, *c*), which indicates the probability that a reference site (on a lipid molecule) has ‘*n*_*d*_’ number of total defect grid points around it within a cutoff radius *r* (here taken to be 14 Å) with ∑χ^2^ value ‘*c*’. These quantities are summed in order to address two important points. The first, is the absence of a one-to-one spatial correlation between the two quantities: χ^2^ is calculated based on lipid coordinates and defect is a grid based analysis that specifically excludes these coordinates. The second, is to incorporate the effect of neighborhood, which can be a determinant factor in mixed phase membrane. In Fig. 3, we present the (*n*_*d*_, *c*) correlation for the three mixed phase membrane systems. For a strictly one-to-one relationship between the two quantities, one can expect data points that are diagonally populated. However, given that dynamics of individual lipids in a mixed phase membrane is delicately connected to its local environment, we can expect a moderately linear relationship with data points smeared around. For a pure phase *L*_*o*_ system, with correlated lipid evolution, the data points in (*n*_*d*_, *c*) correlation are expected to exhibit a narrow range around the diagonal with a further narrow region exhibiting high P(*n*_*d*_, *c*) values near the origin. In contrast, a pure phase *L*_*d*_ system, wherein lipid evolution is not well coordinated, (*n*_*d*_, *c*) correlation can be much wider with an extended region exhibiting high P(*n*_*d*_, *c*) values farther away from the origin along the ∑χ^2^ axis. The profile of (*n*_*d*_, *c*) correlation in a mixed phase membrane can, thus, indicate the global order of the membrane. As observed in Fig. 3, mixed phase DPPC/DOPC/CHOL system exhibit the characteristics of a pure phase *L*_*d*_ system and the mixed phase PSM/POPC/CHOL system resembles a pure phase *L*_*o*_ system. For the PSM/DOPC/CHOL system, with very different χ^2^ distributions for the two pure and mixed membranes (Figure 2), the (*n*_*d*_, *c*) correlation has a wider range (as expected for a pure *L*_*d*_ system), but a narrow region close to the origin exhibiting high probability (as expected for a pure *L*_*o*_ system). Membranes which exhibit a strong correlation, similar to the PSM/POPC/CHOL system here, are therefore more ordered-like and should be dominated by ordered domains. In such a scenario, proteins binding in these systems should also happen in these *L*_*o*_ domains.

**FIG. 3:**
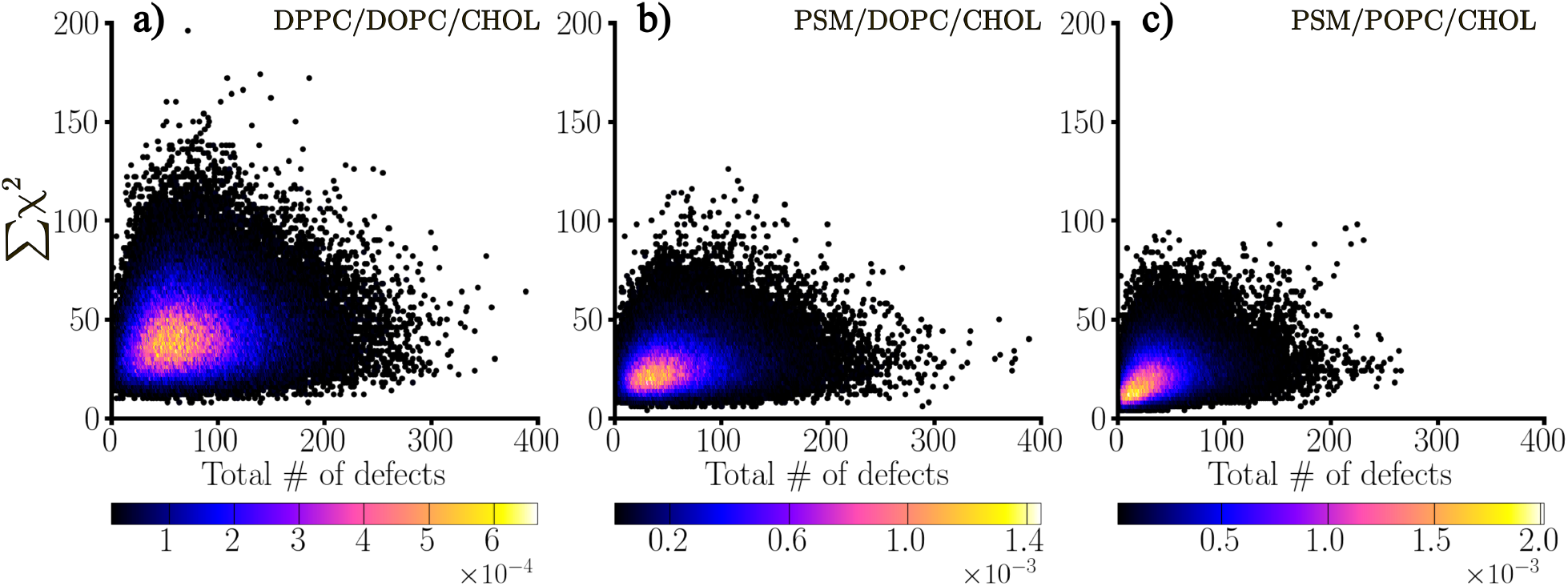
A one-to-one map. χ^2^ and defect correlation map for a) DPPC/DOPC/CHOL, b) PSM/DOPC/CHOL, and c) PSM/POPC/CHOL systems, each exhibiting mixed *L*_*o*_*/L*_*d*_ phase. The colorbars indicate the value of probability P(*n*_*d*_, *c*). The range of axes are kept the same for better comparison.

### Defects in/around *L*_o_ domains exhibit higher persistence than those in *L*_d_ domains

To investigate the functional consequence of such distinct characteristics of mixed phase membranes, we investigate the time evolution of defects therein. Fig. 4 shows the defect spatial maps for the three mixed phase membranes, calculated over 2, 10 and 20 consecutive snapshots, with each snapshot taken at 240 ps time interval. A high probability in the map indicates both the spatial and temporal persistence of a defect, i.e., a defect that is localized on the membrane surface over the analysis window. As expected, the majority of the defects are found to be in the disordered domains (see Fig 1). However, the ordered domains also exhibit a significant amount of defects, including large ones. Moreover, the large defects in (e.g., in PSM/DOPC/CHOL and PSM/POPC/CHOL systems) and around (e.g., in DPPC/DOPC/CHOL) the ordered domains are found to be significantly localized and persist over 20 snapshots, i.e., 4.8 ns (highlighted in Fig. 4 with circles). This is in stark contrast to the defects in the disordered domains which quickly disperse over time. Such ns-long persistent, localized defects in *L*_*o*_ domains is a remarkable observation, since defects are quite transient, owing to the fluidity of the membrane and the thermal fluctuations.

**FIG. 4:**
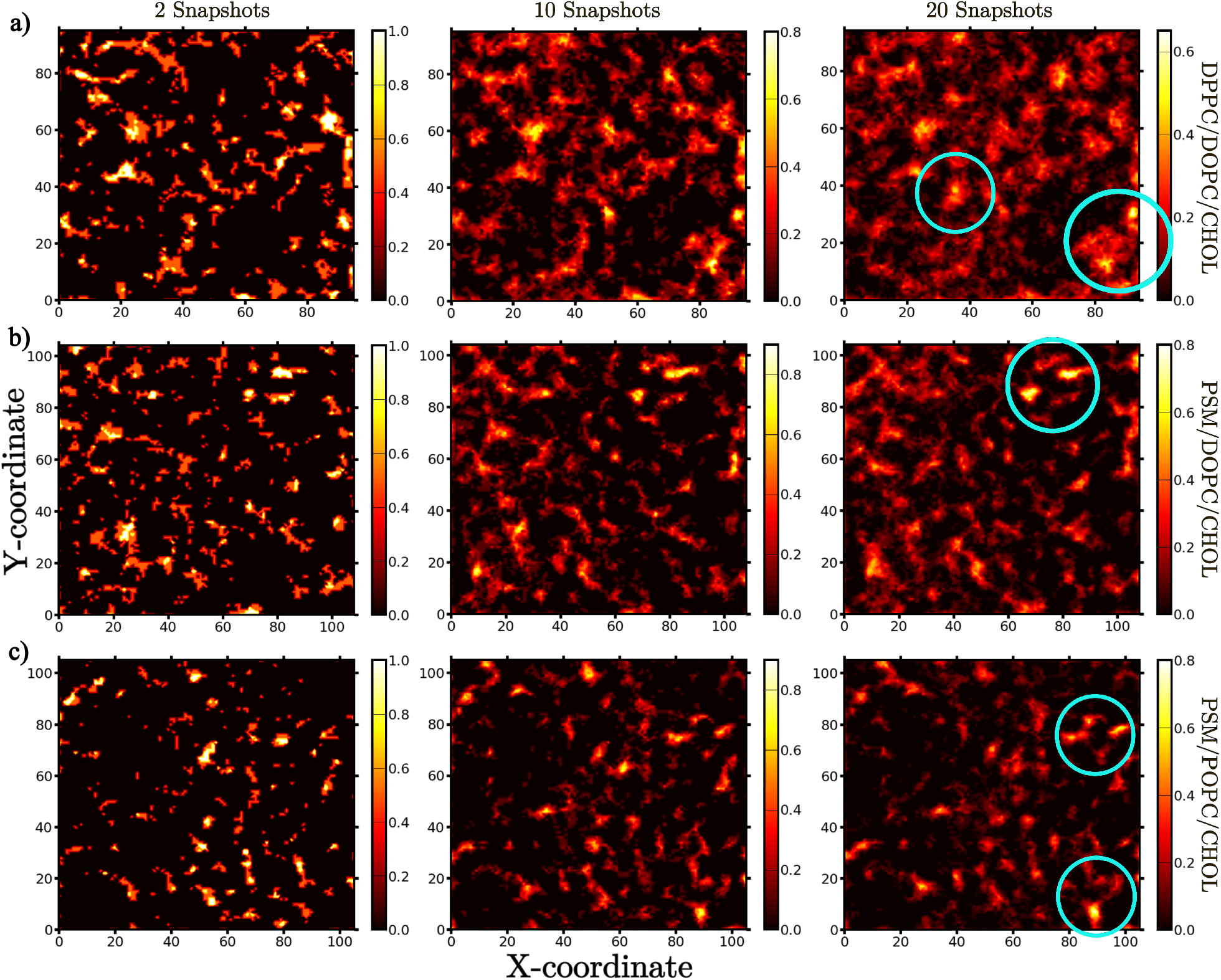
Defect spatial and temporal localization. Defect spatial map of mixed phase a) DPPC/DOPC/CHOL (top row), b) PSM/DOPC/CHOL (middle row), and c) PSM/POPC/CHOL (bottom row) systems calculated over 2 (left column), 10 (middle column), and 20 (right column) snapshots. The cyan circles on the right column indicate the persistent defects in and around the *L*_*o*_ regions.

These observations point us to an important inference: while the disordered domains are rich in large packing defects, the most stable defects exist in and around the ordered domains. This is a consequence of the spatio-temporally correlated evolution of lipids in the ordered domains, which leads to the localization of the packing defects in and around them that can persist over a few nanoseconds. Irrespective of the membrane composition and whether the mixed phased membrane is more ordered- or disordered-like, this feature seems to be a generic one. And likely very important for formation of early encounter sites for peripheral protein binding – especially those that are eventually stabilized due to hydrophobic insertion in the membrane.

### Local membrane order governs the partitioning of membrane-active peptides

Here, we use tLAT as a paradigmatic peptide to explore the molecular origin of peptide localization in a bilayer with phase coexistence and use this example to explore the effect of post transcriptional modification on this localization. Xubo et al.^61^ investigated the role of palmitoylation in the partitioning of tLAT in DPPC/DAPC/Chol (5:3:2) membrane using atomistic and coarse-grained (CG) molecular dynamics (MD) simulation. In both case, they observed the tLAT to partition at the domain boundary, irrespective of the palmitoylation state. Such an observation is in contrast to those reported in experiments, wherein tLAT is shown to partition with *L*_*d*_ domains in GUVs^87^ and *L*_*o*_ domains in GPMVs^86^. While palmitoylation is implicated in the ordered phase affinity^85^, the apparent discrepancy between the observations in GUV and GPMV stem from the relatively strong and weak contrast, respectively, between the raft-like and non-raft domains in the two cases^37^. To further investigate the origin of this apparent discrepancy, Xubo et al.^61^ analyzed the energetics of the lipid-protein interaction in atomistic simulation. Considering DPPC and Cholesterol as *L*_*o*_ domain lipids and DAPC as *L*_*d*_ domain lipids, they computed the residue-wise peptide-lipids interaction energy and observed that palmitoylated tLAT exhibited higher interactions with the *L*_*o*_ domain lipids, while tLAT without palmitoylation interacted strongly with the *L*_*d*_ domain lipids, in line with experiments. Finally, they confirmed the order domain preference of palmitoylated tLAT in CG umbrella sampling simulation of phase separated membrane.

Here we revisit the study by Xubo et al. to investigate the order preference of tLAT, where membrane order is now characterized in terms of the non affine deformation measure i.e, the NAP. Our previous works on both pure phase and mixed phase membranes ^63,88^ have shown that membrane order is dictated by the local lipid packing and their dynamic topological evolution, rather than their specific chemistry. Therefore, there is no a priori reason to assume that all saturated lipids and cholesterol belong to the *L*_*o*_ domain and all unsaturated lipids to the *L*_*d*_ domain. Towards this, we analyzed the atomistic DPPC/DAPC/Chol trajectories used by Xubo et al.^61^. The system consisted of 200 DPPC lipids, 120 DAPC lipids and 80 Cholesterols (200 lipids per leaflet) with a single tLAT with and without palmitoylation. While palmitoylation, in general, is known to enhance the ordered phase affinity of tLAT^37,85^, palmytiolation of a single tLAT can not significantly affect the global membrane order. Nonetheless, the local environment of tLAT in the two cases should bear signature of the order preference of the palmitoyl group. To quantify this, we characterized the two tLAT-membrane systems in terms of the χ^2^ parameter, which, as we have shown earlier, is a sensitive marker of membrane order.

For the χ^2^ analysis, we considered two reference sites for the lipids (C27 and C37 tail carbon atoms for DPPC and DAPC) and one reference site for cholesterol (C9 atom). We considered 4 to 6 reference sites for tLAT with and without palmitoylation that lie close to the lipid reference sites in terms of their *Z* coordinates. Together, we had around 365 reference sites for the C- and N-terminus side of the bilayer in both cases. We computed the χ^2^ values for each reference sites using a neighborhood cutoff Ω = 18 Å (see Method). These values indicate the distinct spatio-temporal evolution of the lipids and tLAT in the membrane. To understand how distinct this topological evolution is for a single tLAT with and without palmitoylation, we need to compare this metric in a local environment around the tLAT. Taking the average coordinate of the 4-6 reference sites of tLAT as the center, we subsequently averaged the χ^2^ values of all lipid and cholesterol reference sites within a radius *r* = 10 Å around it. The resulting quantity ⟨χ^2^⟩_*r*_ provides a quantitative measure of the average orderliness of the local membrane environment around tLAT. The choice of the cutoff radius is motivated by the fact that it should be large enough to include several neighboring lipid reference sites, while small enough to represent the local environment of tLAT. Given that we are interested in the changes in lipid packing around a single tLAT with palmitoylation, such information will be lost with a very large *r*. Finally, we calculated the distribution of ⟨χ^2^⟩_*r*_ values for the two leaflets of the bilayer and subsequently averaged them to get the final probability distribution of ⟨χ^2^⟩_*r*_ for tLAT with and without palmitoylation, which is compared in Fig 5. The distribution of ⟨χ^2^⟩_*r*_ clearly indicates the local environment of palmitoylated tLAT to be relatively more ordered than that of tLAT without palmitoylation. Thus, the non-affine deformation based analysis is sensitive enough to distinguish between the local environment of a single tLAT peptide, without any prior assumption or knowledge of lipid chemistry or membrane composition.

**FIG. 5:**
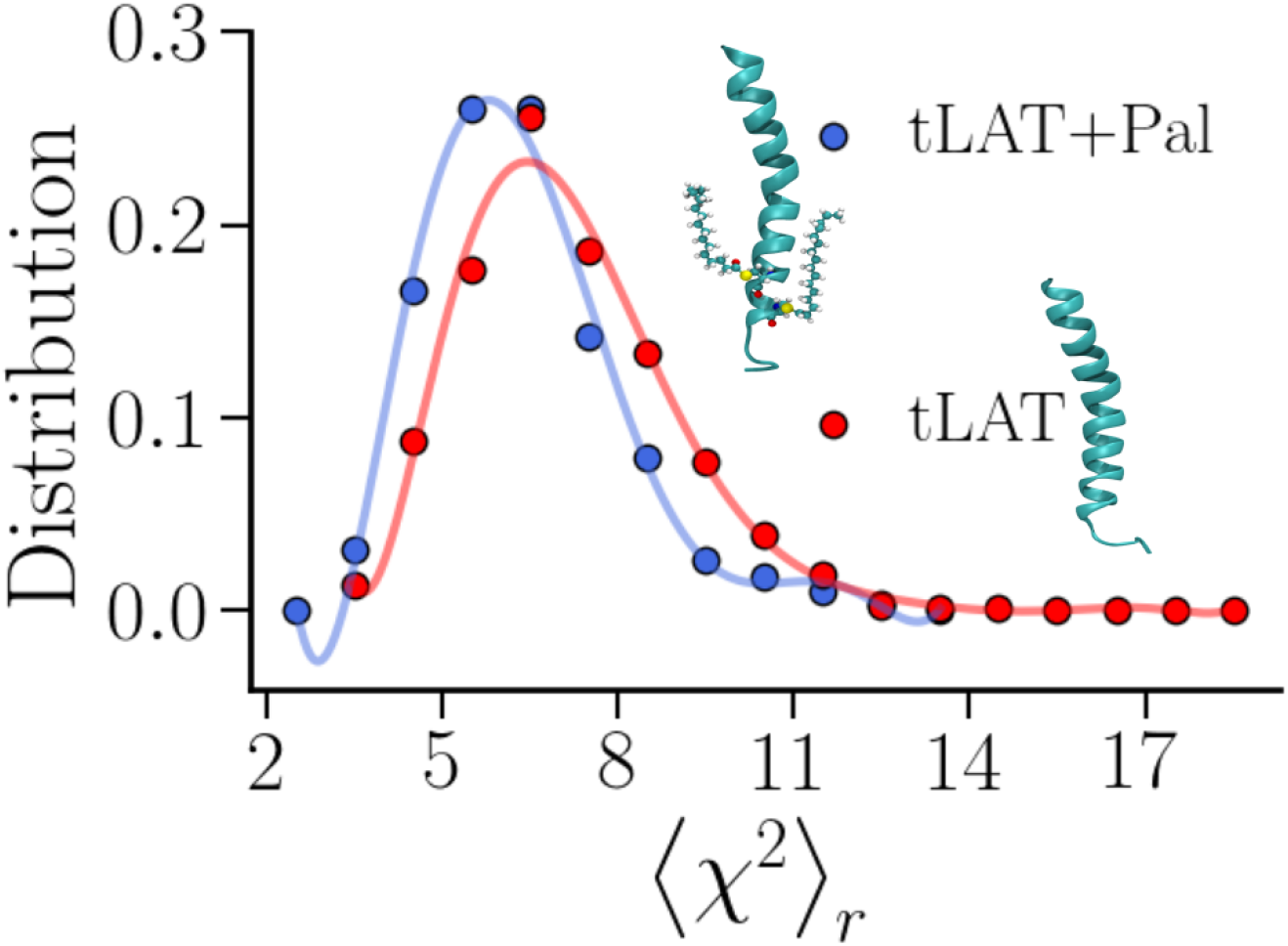
Local order of membrane around tLAT with and without palmitoylation. Distributions of χ^2^ values averaged within a local neighborhood of radius 10 Å around tLAT and palmitoylated tLAT. The lines are polynomial fit to the actual data and are just guides to the eyes.

### Membrane defect profile governs the permeability of small molecules

The defect size distributions for pure phase DPPC/DOPC/CHOL membranes shown in Figure 2 are computed using trajectories that Ghysels et al.^89^ used to analyze water permeation through the membrane and observed a similar permeation mechanism as that of oxygen. As evident, the pure *L*_*d*_ membrane exhibits much larger packing defects than the pure *L*_*o*_ counterpart, which, as discussed earlier, is a generic property of pure phase membranes irrespective of the membrane composition and lipid chemistry. In the context of penetrant permeation through the membrane, these observation would indicate the abundance of permeation channels in the *L*_*d*_ membranes, while the *L*_*o*_ membranes remain sparse. This is in line with the observations from Ghysels et al. that *L*_*d*_ membranes are statistically more permeable. However, the disparity between the lateral and transverse diffusion profile of water/oxygen demand a closer inspection.

To understand this, we further analyzed the oxygen permeation trajectories of Ghysels et al.^89^ (pure *L*_*o*_ and *L*_*d*_ phase DPPC/DOPC/CHOL membranes with O_2_, see Table III) in terms of the packing defects. As in the case of water permeation trajectories, defects on a leaflet in the *L*_*d*_ system were larger in size as compared to the *L*_*o*_ one (Fig 6b). Subsequently, we identified two kinds of defects following our 3-dimensional defect algorithm^71^: surface and core defects. The core defects were identified for membrane slices of thickness 14 and 8 Å around the membrane mid-plane, while the surface defects were identified on one leaflet of the membrane excluding the core region (Fig 6a). The surface and core defects can be interpreted as the free volume available on the membrane surface (in lateral directions) and in its core, respectively, which has long been implicated in lipid diffusion^68,90,92,95^. From the methodology point of view, it is worth mentioning that the defects or free volume in our analysis correspond to the accessible surface areas, which are calculated using a rolling probe like algorithm with a probe radius of 1.4 Å (roughly the radius of a water molecule)^71^. While this is technically justified for the surface defects, which are indeed accessible to water, the free volumes in the core should not necessarily be accessible to water unless they are connected to surface defects. In such a case, using the same probe radius allows for a fair comparison between the two kinds of free volume while providing a conservative estimate of the same. Moreover, the use of periodic boundary condition (along *X* and *Y* directions) is avoided while calculating defect sizes in both cases without the loss of generality to avoid statistical implications of comparing defects in a thin slice of membrane to those on surface. While both these points can lead to under prediction of defect sizes, unintended over predictions (which can also be misleading) can be avoided.

**FIG. 6:**
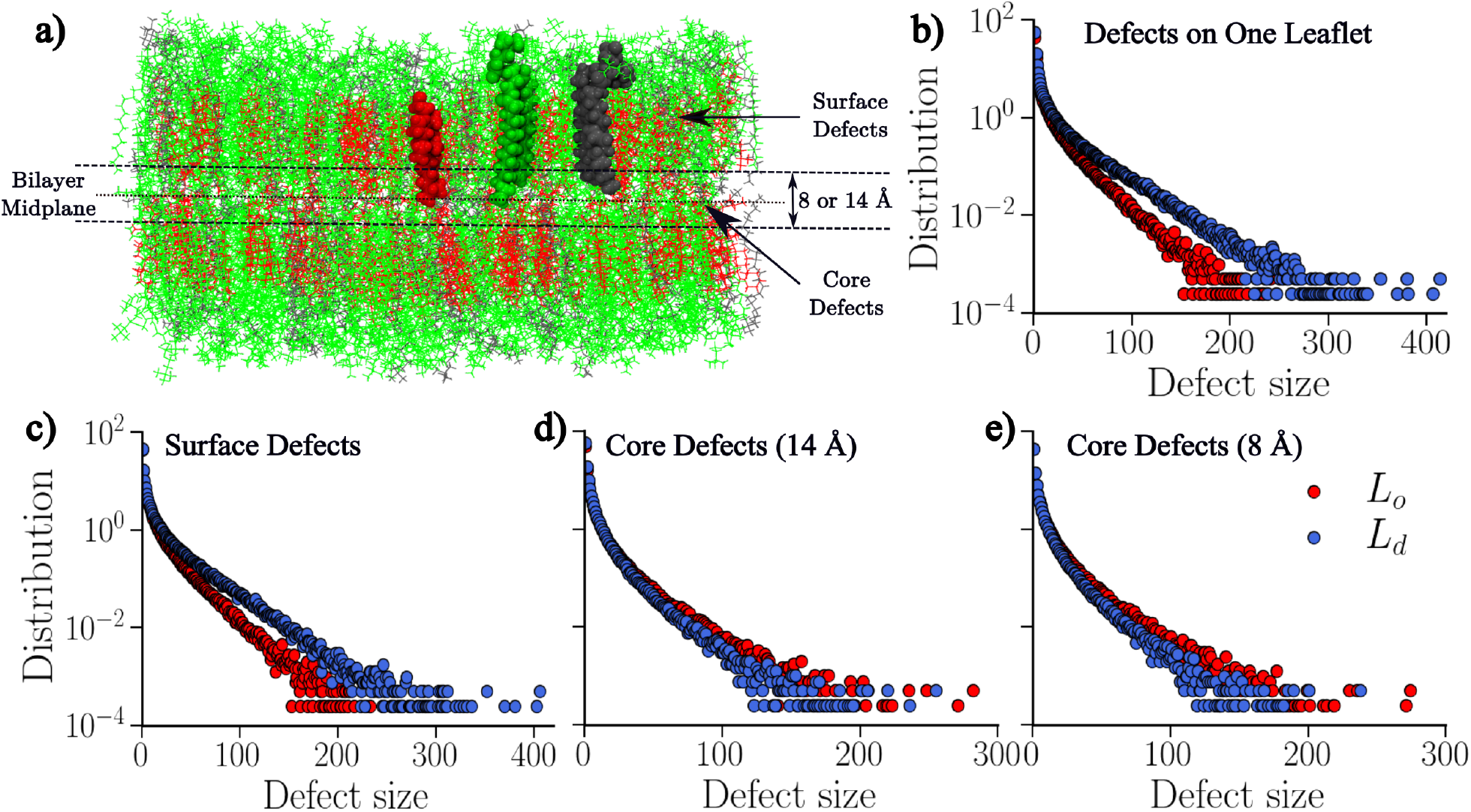
Role of defects/free volume in membrane permeability. a) The definition of various kinds of defects analyzed in the DPPC/DOPC/CHOL membranes containing O_2_ and exhibiting pure *L*_*o*_ and pure *L*_*d*_ phases. One lipid from each kind is shown in van der Waal representation against all other in line representation (DPPC:green, DAPC:grey, CHOL:red). The membrane snapshot has been rendered using the visualization tool VMD^98^. Distributions of defect size for: b) defects on one leaflet, c) surface defects, and core defects within a slab of thickness d) 14 Å and e) 8 Å around the membrane mid-plane.

Fig 6c-e show the size distributions of these defects in the two pure phase membranes. Similar to defects on a single leaflet (Fig 6b), *L*_*d*_ system was found to exhibit larger surface packing defects as compared to the *L*_*o*_ one (Fig 6c), implying more number of permeation channels and stronger transverse diffusion in the *L*_*d*_ system. Together, this can result in significantly larger permeability as compared to *L*_*o*_ system, as reported by the authors. Interestingly, the trend is reversed for the core defects: the *L*_*o*_ system exhibits larger core defects than the *L*_*d*_ system (Fig 6d). Further, as the thickness of the membrane slice is decreased from 14 to 8 Å, the distinction between the size distributions for core defects becomes more apparent (Fig 6e), indicating that the membrane mid-plane of the *L*_*o*_ system has relatively more free volume as compared to the *L*_*d*_ system. This can be attributed to the fact that lipids in the ordered sub-domains in the *L*_*o*_ phase are closely packed in the transverse direction. This circumvents interdigitation of the lipid tails between the leaflets, which can allocate substantial amount of free volume in the mid-plane region. This would provide an ideal path for the penetrants to diffuse along the membrane mid-plane through the isolated free volumes that evolve over time: an observation that provides a possible explain on why the authors observe stronger lateral diffusion in the *L*_*o*_ systems. The trend reversal between surface and core defect size can lead to the anisotropy in the diffusion profile between the two phases, as reported by the authors^89^.

As discussed earlier, there exists a direct correlation between membrane order and packing (here, free volume), irrespective of the membrane composition. It is, therefore, evident that the permeation profile for small molecules in other model membrane should follow the same characteristic: stronger transverse diffusion in *L*_*d*_ membranes vs. stronger lateral diffusion in *L*_*o*_ membranes. This is indeed what Ghysels et al.^89^ observed in terms of free energy profiles and nearest neighbor analysis of water permeation in homogeneous DPPC/DOPC/CHOL, PSM/DOPC/CHOL, and PSM/POPC/CHOL membranes.

## DISCUSSION

It is evident that the relationship between membrane order and defects can be quantified as to be moderately linear. A stricter correlation indicates the mixed phase membrane to be more ordered-like with spatio-temporally correlated lipid evolution and smaller packing defects. In contrast, a moderate correlation indicates the mixed phase membrane to be more disordered-like with uncorrelated lipid evolution and larger packing defects. This correlation can govern various functional aspects of the raft-like domains including protein adsorption and membrane permeation.

While most of the peripheral proteins bind to the *L*_*d*_ domain^99,100^, the specificity of defect profiles in mixed phase membranes suggest that in a membrane that is more ordered-like, binding should also happen in the *L*_*o*_ domain. This is also in line with the experimental observation that some proteins (e.g., tLAT) which exclusively bind to *L*_*d*_ domains in GUVs can also bind to *L*_*o*_ domains in GPMV, wherein the demarcation between the two domains are not too strong^37^.

Furthermore, the distinctive time evolution of defect profiles, as indicated in Fig. 4, can lead to two possible routes for a binding event (see the schematics in Fig. 7). In case I, a protein can sample the membrane surface and bind to a transient, but large defect in the *L*_*d*_ domain. In case II, a protein can bind to a localized, but relatively smaller defect in or around the *L*_*o*_ domain. The two binding routes can be distinguished in terms of a number of features, the first being the time scales of binding, i.e., the time taken by the protein to bind to the membrane. This time scale should be longer in case I as the protein has to extensively sample the membrane surface to bind to large defects that are already diffusing continuously. Accordingly, the strength of the membrane-protein interactions should be stronger in case II than in case I. Finally, the mechanism of defects coalescing upon protein binding should also be different for the two cases. The binding of APLS motifs onto membrane surface proceeds via such coalescence of smaller defects so as to accommodate its hydrophobic surface^70^. Here in case II, defect coalescence can only proceed via modification of the lipid packing upon the protein binding, whereas In case I, thermal fluctuation will suffice given that the defects are already quite transient. While substantial amount of work is needed to establish the existence of these two diverse binding routes, it can certainly elucidate the basic design principles behind membrane-protein partitioning. This in turn can unravel the physiological functions of lipid rafts and also facilitate the design and engineering of proteins that target specific membrane regions.

**FIG. 7:**
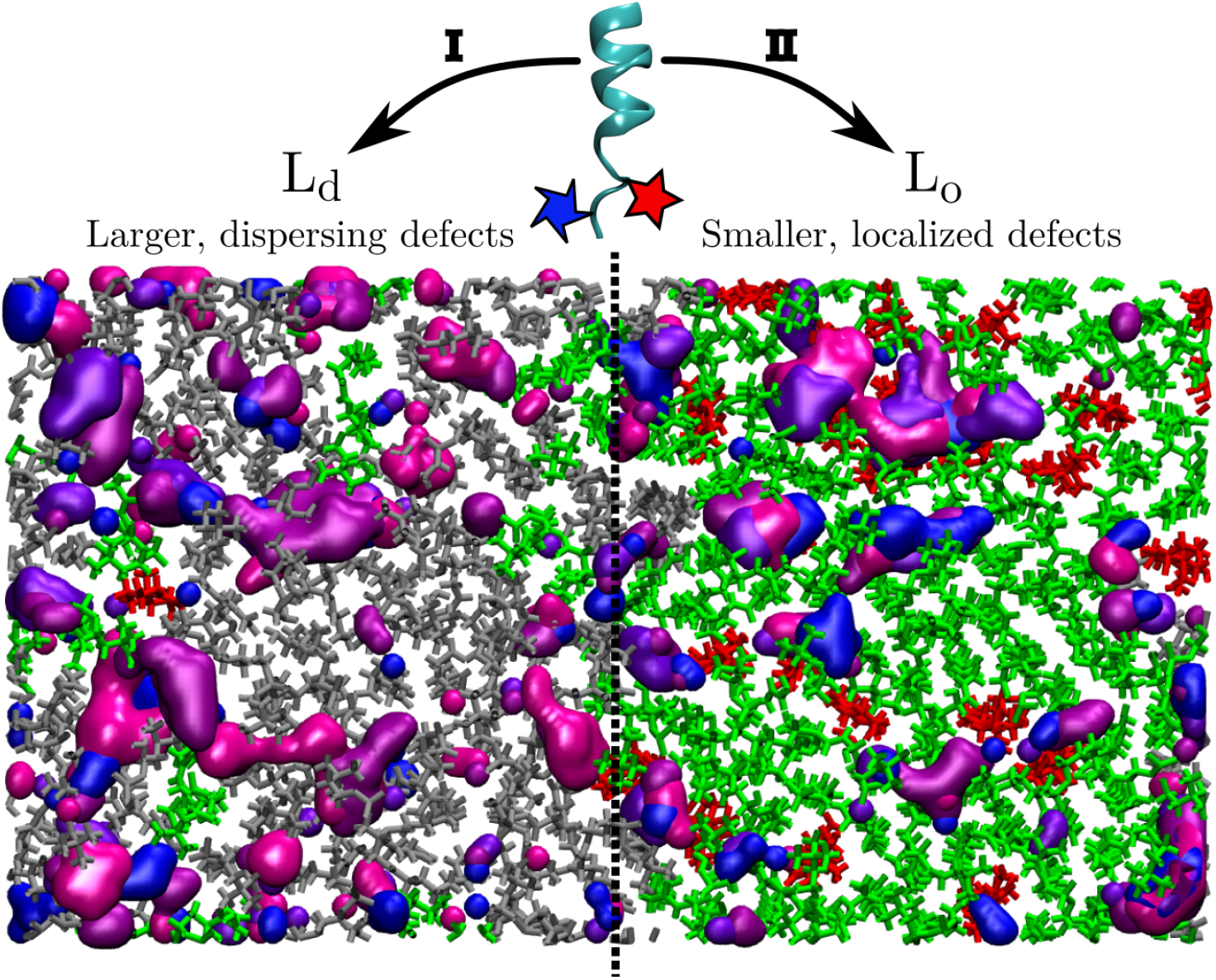
Schematic of defect profile on a lipid membrane exhibiting mixed *L*_*o*_*/L*_*d*_ phase and how it can govern protein partitioning. The top view of membrane is shown with different kinds of lipids in bond representation colored in silver, green, and red and defect pockets analyzed over four consecutive snapshots shown in QuickSurf representation^101^ with colors blue, violet, purple, and magenta. The membrane snapshot has been rendered using the visualization tool VMD^98^. The overlapping defects in *L*_*o*_ domains and near the interface indicate localized defects. A peripheral peptide with bulky side groups (shown in colored polygons) can bind to larger, dispersing defects in the *L*_*d*_ domain (case I) or the smaller, localized defects in and around the *L*_*o*_ domain (case II). See the main text for discussion.

The effect of membrane order on protein partitioning can be further exemplified in case of tLAT, where palmitoylation is shown to enhance the ordered phase affinity of tLAT in agreement with experiments^37,61,85^. It should be noted that unlike CG simulation, wherein there is a clear phase separation between the *L*_*o*_ and *L*_*d*_ domains, the distinction between domains is not so clear in atomistic simulation. This makes the clear characterization of the two phases, in terms of parameters such as area per lipid and bilayer thickness, quite difficult. Accordingly, the analyses on tLAT-lipid energetics by Xubo et al.^61^ relies on the identification of lipids as belonging to *L*_*o*_ or *L*_*d*_ domains based on their chemical identity as saturated lipid (and cholesterol) or unsaturated lipid, respectively. In contrast, the characterization in terms of χ^2^ relies on the information on spatial and temporal evolution of lipids without any information on membrane chemistry and can quantitatively distinguish the subtle difference in the local membrane order around tLAT with and without palmitoylation, even for such a small atomistic system.

Our inference is also in line with the recent observations by Sikdar et al^102^, which relates the impaired membrane activity of hepatitis A virus-2B (HAV-2B) protein in cholesterol rich membranes to the scarcity and small size of defects therein. The ordering effect of cholesterol in lipid membrane is well known^58^ which, in a single component lipid system leads to cholesterol-rich interfacial regions surrounding cholesterol-poor *L*_*o*_ and *L*_*d*_ nanodomains^47^. For a two component lipid mixture, cholesterol preferentially partitions with unsaturated lipids, leading to ordered sub-domains made of of saturated lipids^59,62^. The correlation between membrane order and defects, thus implies significant modification in packing defects in the membrane, which for a more ordered-like membrane results in fewer, smaller and shallower packing defects. Accordingly, presence of cholesterol in membrane can significantly reduce the binding events.

Beyond influencing protein partitioning, membrane packing and order can also influence the permeability of the membrane against small molecules. This can be attributed to the free volume available on the membrane surface and in its core that provides suitable hopping paths to the penetrants. The correspondence between free volume and the permeability of the membrane is rather intuitive and was also mentioned by Ghysels et al.^89^. Herein, we have been able to systematically correlate membrane order and permeability in terms of the packing defects. The membrane can be visualized as made up of ordered regions that act as platforms for strong transverse diffusion owing to the large free volume at the membrane core. The disordered and boundary (between the ordered/disordered domains) regions act as channels for penetrant permeation in/out of the membrane, due to the abundant surface defects that can initialize permeation.

To summarize, we observe some generic trends in the packing defects profiles in mixed phase membranes, which correlate almost linearly with the membrane order and exhibit distinct temporal evolution. The local membrane order crucially governs the preferential partitioning of peripheral proteins, while the membrane defect/free volume governs the membrane permeability of small molecules. The specificity of defects in mixed phase membranes can, therefore, have important lateral and trans-bilayer functional implication and might also follow the same basic design principles based on membrane order and packing.

## MATERIALS AND METHODS

### A. Details of the simulation trajectories

The pure and mixed phase atomistic trajectories of model ternary lipid systems (total 9), analyzed in the first part of the work, were obtained from Edward Lyman’s group at the University of Delaware^62^. The trajectories were equilibrated in NAMD^103^ using TIP3P water model ^104^, CHARMM36 lipid force field ^105^, and Pitman et al.’s^106^ cholesterol model. The *μs* long simulations were performed in Anton supercomputer^107,108^. The relevant details on system composition and size are summarize in Table I. Readers are directed to the original references^59,62^ for further technical details on the system preparation and simulation methodologies.

**TABLE I:** Details of the atomistic trajectories of pure and mixed phase systems used to analyze the correlation between membrane order and packing

| Membrane system | Membrane Phase | Lipid Composition | Total # Lipids | System size ($\sim \text{nm} \times \text{nm}$) | Trajectory Length ($\mu s$) |
| --- | --- | --- | --- | --- | --- |
| DPPC/DOPC/CHOL | $L_o$ | 0.55/0.15/0.30 | 512 | $10.4 \times 10.4$ | 5.05 |
| | $L_d$ | 0.29/0.60/0.11 | 400 | $10.8 \times 10.8$ | 2.02 |
| | $L_o/L_d$ | 0.37/0.36/0.27 | 380 | $9.55 \times 9.55$ | 9.43 |
| PSM/DOPC/CHOL | $L_o$ | 0.64/0.03/0.33 | 560 | $10.62 \times 10.62$ | 0.95 |
| | $L_d$ | 0.15/0.82/0.03 | 574 | $13.44 \times 13.44$ | 0.81 |
| | $L_o/L_d$ | 0.43/0.38/0.19 | 452 | $10.86 \times 10.86$ | 4.54 |
| PSM/POPC/CHOL | $L_o$ | 0.61/0.08/0.31 | 560 | $10.65 \times 10.65$ | 9.08 |
| | $L_d$ | 0.23/0.69/0.08 | 560 | $12.4 \times 12.4$ | 1.14 |
| | $L_o/L_d$ | 0.47/0.32/0.21 | 506 | $10.56 \times 10.56$ | 6.88 |

The all-atom trajectories for tLAT-lipid membrane system with and without the palmitoyl group were obtained from Xubo et. al.^61^. The relevant system details are provided in the main text and are summarized in Table II. The oxygen permeation trajectories were borrowed from the study of Ghysels et al.^89^. The system consisted of 50 oxygen molecules in DPPC/DOPC/CHOL membrane systems exhibiting pure *L*_*o*_ and pure *L*_*d*_ phases, whose details are summarized in Table III. Further technical details on the above systems can be found in the original references^61,89^.

**TABLE II:**
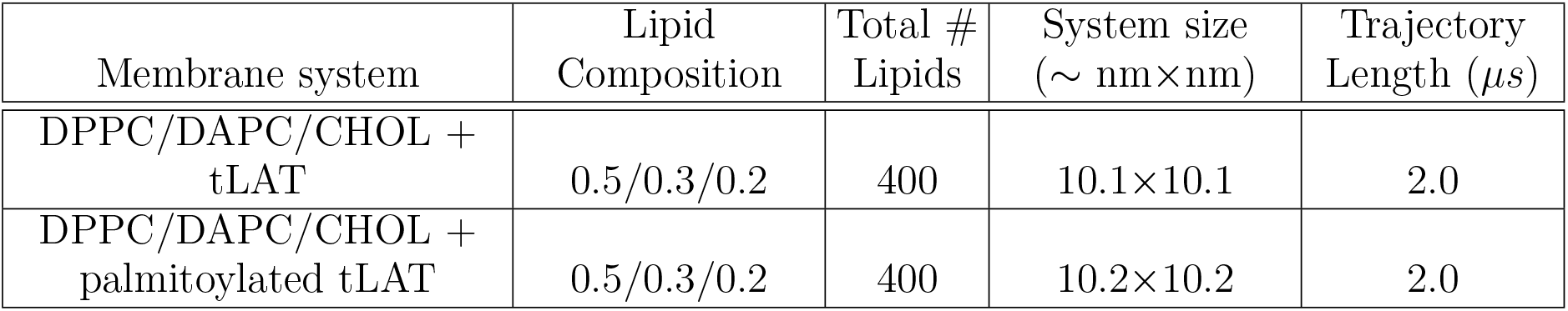
Details of the atomistic trajectories used to analyze partitioning of tLAT with and without palmitoylation

**TABLE III:** Details of the atomistic trajectories used to analyze oxygen permeation in pure phase DPPC/DOPC/CHOL membranes

| Membrane system | Membrane Phase | Lipid Composition | Total # Lipids | System size (nm×nm) | Trajectory Length ($\mu s$) |
| --- | --- | --- | --- | --- | --- |
| DPPC/DOPC/CHOL + 50 O<sub>2</sub> | pure $L_o$ | 0.55/0.15/0.30 | 512 | 10.88×10.88 | 1.14 |
| | pure $L_d$ | 0.30/0.62/0.08 | 400 | 11.95×11.95 | 0.95 |

### B. Calculation of the non-affine parameter (NAP) (χ^2^)

The χ^2^ analysis is performed as described in our previous studies^63,88^ and follows the original prescription by Falk and Langer as applied for amorphous solids^96^. The lipid membrane is considered as an amorphous system evolving in space and time, where a reference site on every lipid molecule is tracked over time. In our case, unless otherwise noted, the reference sites are identified as the bottom carbon atom of the glycerol group for lipids and the hydroxyl group oxygen atom for cholesterol. A neighborhood is defined around each central lipid within a cutoff radius Ω (see Fig. 1a), which is taken to be 14 Å in our calculations (unless otherwise specified). As discussed previously^63,88^, χ^2^ is calculated at each reference site in the membrane as

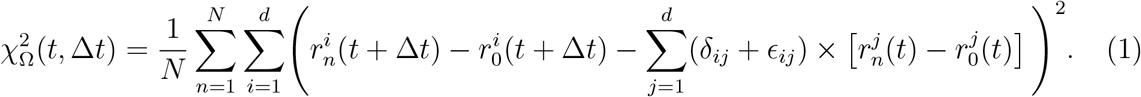

where *i, j* indicate spatial coordinates *r* of the lipid reference site with dimension *d, n* runs through the *N* lipids in the neighbourhood Ω around the central lipid *n* = 0 (Fig. 1a), and *δ*_*ij*_ denotes the Kroneker delta function. *ϵ*_*ij*_ which denotes the strain associated with the maximum possible affine part of the deformation, minimizes 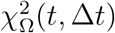 and is calculated as follows

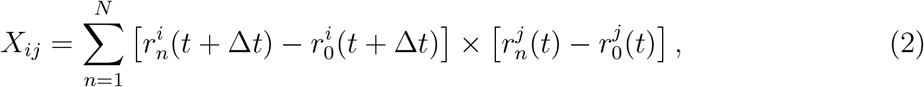

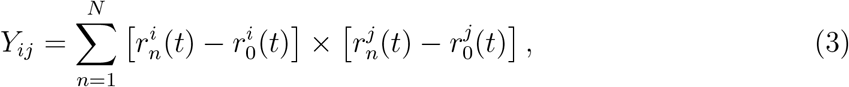

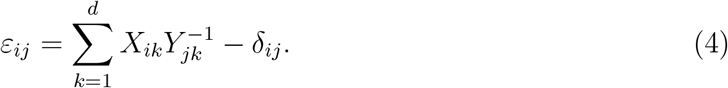

As opposed to the previous implementations^63,88,96^, we now normalize the χ^2^ by the total number of lipids in the neighbourhood *N*, so as to incorporate the effect of environment in mixed phase membranes and also to be able to compare the results across various phases (*L*_*o*_, *L*_*d*_, or mixed) and systems of various sizes (see Table I).

To generate the χ^2^ spatial maps, we average the computed χ^2^ values, over 10 consecutive system snapshots and map that to the spatial coordinates of the reference sites of lipids of the middle (sixth) snapshot of the chosen set (Fig. 1c).

The code to calculate the NAP is available on the Github repository https://github.com/codesrivastavalab/DegeneracyBiologicalMembraneNanodomains/tree/master/codes-degeneracy-membrane-nanodomains^27^.

### C. Identifying and analyzing three dimensional packing defects

We identify the packing defects/free volume in lipid membranes based on our three dimensional defect algorithm^71^, which is based on a grid based free volume analysis. The defects are identified as grid points that lie around the exposed hydrophobic tails of the lipids and below the lipid reference sites (the bottom carbon atom of the glycerol group for lipids and the hydroxyl group oxygen atom for cholesterol). For the pure and mixed phase membrane systems under study, we identify the defects on one leaflet based on a suitable Z-cutoff. The defect grid points are subsequently grouped following a distance based clustering approach to identify the individual defect pockets and their size distributions are calculated.

The defect spatial maps (Fig. 1d, Fig. 4) are calculated by projecting the defects at each snapshot onto the XY grid plane, binning them (bin size: 1 Å) and finally averaging over a set of snapshots. The resulting count indicates the probability that a grid point belongs to a defect and a higher probability indicates both the spatial localization and temporal persistence of the defect.

To corroborate the findings of Ghysels et al.^89^ in terms of packing defects, we identify two more classes of defects. The surface defects basically are the hydrophobic defect pockets on membrane surface (same as earlier), which are identified on a membrane leaflet with Z-coordinates *>* 10 Å. The core defects indicate the free volume in the membrane mid-plane and are identified as the grid points that are not occupied by the lipids/cholesterol. For both classes of defects, we use a probe radius of 1.4 Å to identify defect grid points (as in a rolling-probe method, see the original reference for an in-depth discussion^71^). We do not include periodic boundary condition while calculating the defect size distribution. This allows us to compare both classes of defects without any loss of generality.

The code to identify 3-dimensional packing defects is available on the Github repository https://github.com/codesrivastavalab/3DLipidPackingDefects^71^.

## AUTHOR CONTRIBUTIONS

MT and AS designed the research. MT performed the research and analyzed the data with AS. MT wrote the paper with help from AS.

## ACKNOWLEDGEMENT

The authors thank Edward Lyman (University of Delaware) for sharing the simulation trajectories and acknowledge Anton supercomputer facility for making the trajectories available. The authors thank Xubo Lin and Alemayehu A. Gorfe (The University of Texas Health Science Center at Houston, Texas) for sharing the tLAT trajectories, and Richard M. Venable and Richard W. Pastor (Laboratory of Computational Biology, National Heart Lung Blood Institute, NIH) for sharing the oxygen diffusion trajectories.

MT would like to acknowledge financial support from DBT and IISc-Bangalore. Financial support from the Indian Institute of Science-Bangalore and the high-performance computing facility “Beagle” setup from grants by a partnership between the Department of Biotechnology of India and the Indian Institute of Science (IISc-DBT partnership programme) are greatly acknowledged. AS thanks the startup grant provided by the Ministry of Human Resource Development of India and the early career grant from the Department of Science and Technology of India. AS also thanks the DST for the National Supercomputing Mission grant (DST/NSM/RD-HPC-Applications/2021/03.10). FIST program sponsored by the Department of Science and Technology and UGC, Centre for Advanced Studies and Ministry of Human Resource Development, India is gratefully acknowledged by the authors. This research was also supported in part by the National Science Foundation under Grant No. NSF PHY-1748958 (KITP e-visit).

## Notes

### Competing Interest Statement

The authors have declared no competing interest.

